# Early-Stage Identification and Avoidance of Antisense Oligonucleotides Causing Species-Specific Inflammatory Responses in Human Volunteers Peripheral Blood Mononuclear Cells

**DOI:** 10.1101/2021.10.30.466173

**Authors:** Sebastien A. Burel, Todd Machemer, Brenda F. Baker, T. Jesse Kwoh, Suzanne Paz, Husam Younis, Scott P. Henry

## Abstract

A human peripheral blood mononuclear cell (PBMC)-based assay was developed to identify antisense oligonucleotide (ASO) with the potential to activate a cellular innate immune response outside of an acceptable level. The development of this assay was initiated when ISIS 353512 targeting the mRNA for human C-reactive protein was tested in a phase I clinical trial in which healthy human volunteers unexpectedly experienced increases in interleukin-6 (IL-6) and C-reactive protein. This level of immune stimulation was not anticipated following rodent and non-human primate safety studies in which no evidence of exaggerated proinflammatory effects were observed. The IL-6 increase induced by ISIS 353512 was caused by activation of B-cells. The IL-6 induction was inhibited by chloroquine pretreatment of the PBMC and the nature of the ASOs suggested that the response is mediated by a toll like receptor, in all likelihood TLR9. While assessing the inter PBMC donor variability, two class of responders to ISIS 353512 were identified (discriminator and non-discriminators). The discriminator source of PBMC was shown to produce low level of IL-6 after 24 hours in culture in absence of ASO treatment. The PBMC assay using discriminator donors was shown to be reproducible allowing to assess reliably the immune potential of ASOs via comparison to known benchmark ASO controls that were previously shown to be either safe or inflammatory in clinical trials.

## Introduction

Antisense oligonucleotides (ASO) are short, synthetic single strands of nucleic acids with phosphorothioate linkage that hybridize to messenger ribonucleic acid (mRNA) through Watson-Crick base pairing (Stephenson and Zamecnik 1978). They are designed to alter the expression of a protein by selectively binding to the RNA that encodes the targeted protein (Crooke 2004). In recent years, ASOs have been used as therapeutic agents to inhibit the expression of a specific target mRNA transcript involved in a growing range to diseases (Crooke, Witztum et al. 2018), several of them, Waylivra (volanesorsen), Kynamro (mipomersen) and Tegsedi (inotersen), have received regulatory acceptance for commercialization (Hovingh, Besseling et al. 2013, Benson, Waddington-Cruz et al. 2018, Gouni-Berthold, Alexander et al. 2021). However, despite ASO being generally well tolerated by human volunteers and patients (Crooke, Baker et al. 2016), some phosphorothioate containing ASOs have been shown to elicit nonspecific and proinflammatory effects of varying intensity in a sequence-specific manner (Monteith, Henry et al. 1997). A number of modifications of the ASO designs have been introduced such as incorporation of 2’-O-methoxyethyl (MOE) and systematic methylation of cytosine to mitigate these proinflammatory effects. Despite those improvements, a small fraction of sequences examined in rodents have exhibited a higher level of proinflammatory effects (Paz, Hsiao et al. 2017). Those proinflammatory effects have been linked to a growing network of receptors and coreceptor involved in the recognition of both DNA and RNA oligonucleotides (ODNs) (Senn, Burel et al. 2005, Sioud 2010, Burel, Machemer et al. 2012, Meng and Lu 2017, Paz, Hsiao et al. 2017), including one family of the cell surface and endosome-associated receptor, toll-like receptors (TLRs), which has been shown to recognize synthetic nucleic acids (Krieg, Yi et al. 1995, Krieg 1999, Akira and Takeda 2004). One of the members of this family, TLR9, can recognize unmethylated cytosine-guanosine (CpG) motifs which might be present in oligonucleotides (ODNs) and mediates of proinflammatory effects (Jurk, Schulte et al. 2004, Abel, Wang et al. 2005, Marshall, Fearon et al. 2005). These unmethylated CpG motifs, common in bacteria and DNA viruses, are mostly suppressed and methylated in vertebrate genomes. While some therapeutic ODNs incorporate CpG motifs to elicit therapeutically beneficial proinflammatory effects for a range of indications (Campbell 2017, Karapetyan, Luke et al. 2020), most therapeutic ASOs have been optimized to minimize risk of proinflammatory response by methylating every cytosine in the ODN sequence as well as avoiding the presence of CpG motifs.

Historically, the bulk of the screening experience with ASOs has been gained in rodents. Overall, rodents have been shown to be the most sensitive species, exhibiting a range of inflammatory features ranging from elevated chemokines, increased organ weight attributed to lymphohistiocytic infiltration and various inflammatory cell proliferation (Younis, Vickers et al. 2006, Frazier 2015). As a result of this rigorous screening process, the incidence of proinflammatory effects observed in the patients such as injection site reactions and constitutional symptoms has decreased resulting in better tolerated ODN in a clinical setting.

In this study, we report the case of a 2’MOE ASO, ISIS 353512 which was developed originally to inhibit human C-reactive protein (hCRP) mRNA diseases and as a result reduce level of human CRP in the serum (hsCRP) to decrease the severity of cardiovascular (Pepys, Hirschfield et al. 2006, Szalai, McCrory et al. 2014). In preclinical studies, ISIS 353512 did not reveal any evidence of exaggerated proinflammatory effects in either rodent or non-human primate safety studies. To assess the safety in humans, ISIS 353512 was administered initially to healthy human volunteers in a single and multiple ascending dose double blind randomized trial (NCT00734240). Since ISIS 353512 was intended to reduce levels of human CRP mRNA, serum level of hsCRP were closely monitored during the clinical trial. Unexpectedly caused a higher degree of injection site inflammation, transient constitutional symptoms such as fever, chills, asthenia, feeling hot and feeling cold,… shortly after the first dose and increases in several proinflammatory endpoints such as IL-6 than expected. In particular, hsCRP was markedly elevated in the majority of volunteers treated with a single 100 mg and 200mg dose of ISIS 353512. As a result, clinical investigations of ISIS 353512 were discontinued.

In this manuscript, we present an in vitro method using human peripheral blood mononuclear cells (PBMC) to identify oligonucleotides, such as ISIS 353512, capable of eliciting a human specific proinflammatory response and exclude as a clinical development candidate thereby minimizing the risk of unforeseen inflammatory related adverse events in volunteers or patients. As with any recent ASOs tested in patients, ISIS 353512 did not contain any problematic sequences such as proinflammatory CpG motifs and G quartets. Using hPBMC, we demonstrated that despite the lack of CpG motifs, ISIS 353512 is likely eliciting a human TLR9 mediated proinflammatory response. In hindsight, this outcome is not completely surprising as known differences between rodents and humans in term of the cellular distribution of TLRs and their propensity to recognize slightly different canonical CpG motifs in species dependent fashions has been reported previously (Roberts, Sweet et al. 2005).

## Materials and Methods

### Primate Source and Care

Male and female cynomolgus monkeys were obtained from Guangxi Grandforest scientific primate company, Ltd. (Guangxi, China). All animal procedures were conducted at the Korean Institute of Toxicology (Daejon, Korea) utilizing protocols and methods approved by the Institutional Animal Care and Use Committee (IACUC) and were in compliance with Animal Welfare Act and Guide for the Care and Use of Laboratory Animals (by ILAR publication).

### GLP toxicology and toxicokinetic study of ISIS 353512 in cynomolgus monkeys

Monkeys received either SC injection of saline (vehicle control) or ISIS 353512 at dose levels of 3, 8, 20, and 40 mg/kg/dose on days 1, 8, and 11 (loading doses) and then once weekly for a total duration of 13 weeks, with a 13-week recovery period (saline control, 20 mg/kg and 40 mg/kg groups only). Serum samples were collected at predose, 2, 6, 24, and 48 h following the first dose (Day 1) and last dose (Day 85). Serum samples were analyzed for C-Reactive Protein (CRP) concentration by an ELISA method using IMUCLONE CRP (hs) ELISA kit (American Diagnostica, Stamford, CT).

### ODN design, synthesis and preparation

All oligonucleotides were designed and synthesized at Ionis Pharmaceuticals (supplemental table 1). To identify mouse ASO inhibitors, rapid throughput screens were performed in vitro as previously described (Watts, Manchem et al. 2005). The first 3 to 5 bases and last 3 to 5 bases of chimeric ASO have a 2’-O-(2-methoxy)-ethyl modification, and the ASOs also have a phosphorothioate backbone. ASOs were diluted in phosphate-buffered saline (PBS) for both in vivo and in vitro usage.

### In vivo treatment in mice with ODN

Compounds were dissolved in PBS at varying concentrations as indicated in the results section and figure legends (ie, 150 mg/kg), filtered, sterilized, and administered at 10 mL/g animal weight. Immediately before sacrifice, mice were anesthetized with isoflurane and terminal bleed was performed by cardiac puncture in compliance with Animal Welfare Act and Guide for the Care and Use of Laboratory Animals (by ILAR publication). Plasma was isolated from whole blood and analyzed for clinical chemistries and cytokine levels.

### Human and Cynomolgus peripheral blood mononuclear cells (PBMC) assay

Whole blood was collected from human volunteer donors with informed consent in 8 to 10 BD Vacutainer CPT 8 ml tubes (BD Biosciences, San Jose, CA). The blood samples were mixed immediately prior to centrifugation by gently inverting tubes 8-10 times. The CPT tubes were centrifuged at room temperature in a horizontal (swing-out) rotor for 30 min. at 1500-1800 RCF with brake off. The PBMC were retrieved at the interface between Ficoll and polymer gel layers and transfer to a sterile 50 ml conical tube; pooling up to 5 CPT tubes / 50 ml conical tube / donor. The cells were washed twice with PBS (Ca^++^, Mg^++^ free; GIBCO) prior to another round of centrifugation. The supernatant was discarded without disturbing pellet. The cells were resuspended in RPMI+10% FBS+ penicillin /streptomycin. The cell density was estimated, and the density adjusted as to plate cells at 5 x 10^5^ in 50 μl / well of 96-well, sterile, round bottom, polypropylene plates for each ODN concentration tested for each donor. The ODNs 1:5 serial dilutions [2x] prepared in medium (RPMI+10% FBS+ pen/strep) in a separate sterile, V-bottom, polypropylene plate starting with the highest concentration (400 μM) in the top row.

Added 50 μl / well of 2x concentrated ODN diluted in RPMI+10% FBS+penicillin /streptomycin. The plated cells were incubated for 24 hours at 37°C; 5% CO_2_. Plates centrifuged at 330 x g for 5 minutes before removing supernatants to a separate sterile polypropylene 96-well plate. Cell supernatants stored at −70°C until processing for cytokine assay profiling.

### Magnetic cell sorting of B cells

Human B cells were isolated from PBMC collected from human volunteer donors by magnetic-activated cell sorting using the B cell Isolation Kit II for human cells from Miltenyi Biotec Inc (Auburn, CA) according to the manufacturer’s instruction. Using this approach, B cells are left untouched and ready for subsequent analysis. Human B cells from PBMC were labelled with B Cell lineage-specific CD19 antibodies and removed by magnetic-activated cell sorting by using the Lineage Cell Depletion Kit for human cells from Miltenyi Biotec Inc (Auburn, CA) according to the manufacturer’s instruction. Using this approach, remaining cells (CD19 negative) are left untouched and ready for subsequent analysis. The success or the enrichment or depletion was confirmed by flow cytometry.

### Meso Scale Discovery platform

Multiplex plates pre-coated with capture antibodies for specific cytokines, along with diluents were brought to room temperature prior to use. Diluted standards (3 replicates) and undiluted samples were added to the wells and incubated for two hours in the Meso Scale Discovery^®^ Custom array. All incubations were performed at room temperature with vigorous shaking (300-1000 rpm). After the two-hour incubation, the plate was decanted and washed three times before adding a cocktail of Sulfo-Tag^®^ detection antibodies to each well. After incubating with detection antibodies for two hours, the plate was again decanted and washed three times. Read buffer was added to the wells using reverse pipetting techniques to ensure bubbles are not created to interfere with imaging. The plate was immediately imaged using the MSD Sector 2400 imaging system, and data was analyzed using MSD Discovery Workbench^®^ software. Concentrations of all unknown samples were back-calculated using results interpolated from the corresponding standard curve regression using a weighted, four-parameter fit. Final sample concentrations (pg/ml) were calculated by factoring the dilution factor used for each sample.

### Statistical analysis

The data of in vivo studies (at least four animals per group) are expressed as the mean – standard deviation and were analyzed by ANOVA. When significance was obtained, multiple comparison analysis was conducted using Bonferroni as a post hoc test. In some instances, unpaired Student’s t-test was utilized. All graphs and statistical analysis were obtained using JMP 15 (SAS Institute Inc, Cary, NC).

## Results

### Evaluation of the sub-chronic tolerability of ISIS 353512 in non-human primates and rodents

In order to ensure that risk of adverse events in human healthy volunteers or patients are minimized upon the initiation of the Phase 1 clinical trial, ISIS 353512 was tested for tolerability in cynomolgus monkeys as part of a 3 month repeat dosing toxicology study. ISIS 353512 was administered weekly at 8 and 20 mg/kg/dose (intravenous bolus injection) and 40 mg/kg/dose (subcutaneous injection) for 85 days which revealed no evidence of marked proinflammatory effects. There was neither acute nor sustained induction of serum CRP levels immediately after either a single or multiple dose (figure 1). There were neither acute nor sustained significant alterations in inflammatory cytokines (IL-1β, IL-6, IL-10, IFN-γ, and/or TNF-α) or chemokines (MIP-1α and/or MIP-1β) measured induced by ISIS 353512 except highly variable MCP-1 values at doses ≥ 20 mg/kg/week 2 to 6 hours after injection (data not shown). As with cynomolgus monkeys, repeat administration of ISIS 353512 to Sprague-Dawley rats (13 weeks, 40 mg/kg weekly subcutaneous dosing) did not reveal any adverse effect in any proinflammatory endpoints (data not shown). Therefore, combined findings from non-human primates and rodent studies indicated that administration of ISIS 353512 was tolerated well enough to proceed into human testing and would be unlikely to induce adverse proinflammatory responses in human volunteers.

**Figure 1:**
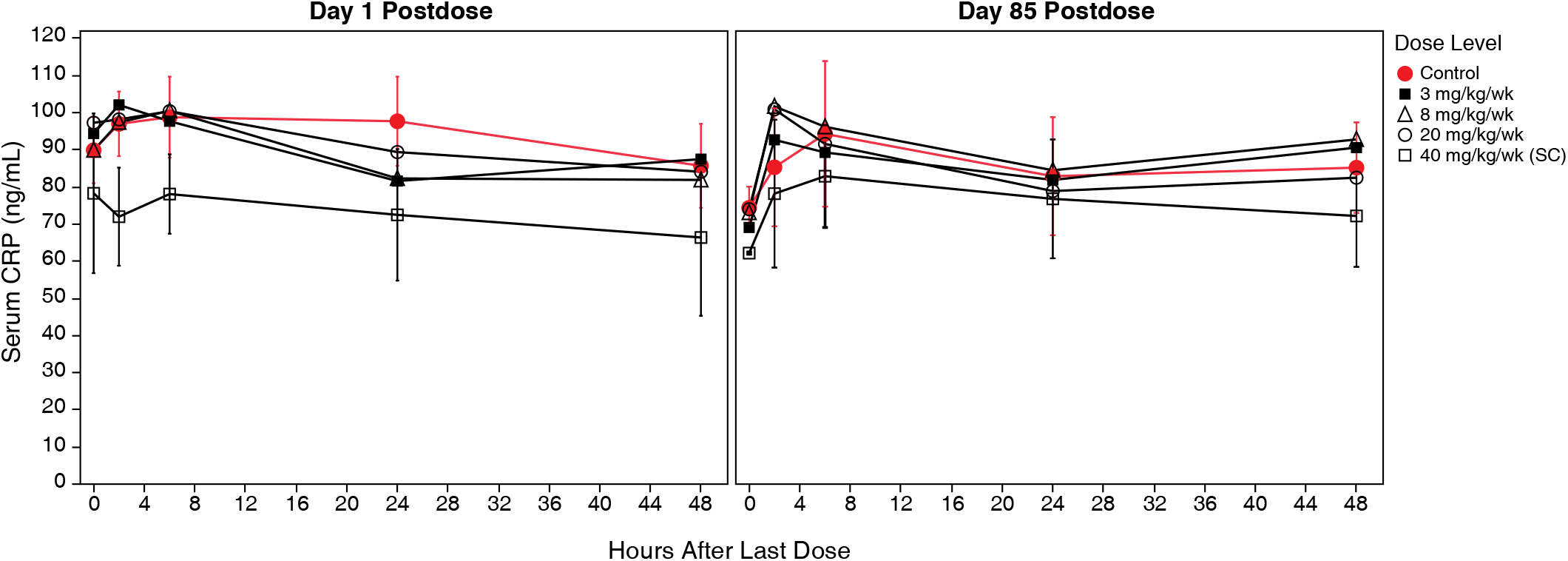
Cynomolgus monkeys were administered ISIS 353512 by IV dosing for control, 3, 8 and 20 mg/kg/week groups and SC dosing for 40 mg/kg/week group. Serum samples were collected at 2, 6, 24 and 48 hours after a single and last dose on Days 1 and 85 and CRP values measured. Means and standard deviations of the CRP responses to ISIS 353512 treatment are shown.

### Development of human PBMC assay to identify proinflammatory oligonucleotides

The discrepancy between the absence of proinflammatory signal in both nonclinical species and the constellation of proinflammatory signals such as elevated CRP or prevalent constitutional symptoms observed in ISIS 353512 Phase 1 trial emphasized the need for an assay able to assess human specific proinflammatory potential of novel ASOs complementing the preclinical toxicology studies.

Despite the lack of canonical CpG motif present on ISIS 353512, we hypothesized that the proinflammatory response observed in human volunteers might be, at least in part, mediated through TLR9 (Krieg, Yi et al. 1995, Bauer, Kirschning et al. 2001, Krug, Rothenfusser et al. 2001, Jurk, Schulte et al. 2004, Abel, Wang et al. 2005, Marshall, Fearon et al. 2005). If indeed ISIS 353512 was able to activate TLR9 signaling, the expression of TLR9 which is mostly limited to B cells and dendritic cells in humans (Krieg, Yi et al. 1995, Krug, Towarowski et al. 2001, Peng 2005), suggested that human peripheral blood mononuclear cells (hPBMC) might provide a suitable model to assess the proinflammatory potential of ISIS 353512. Peng et al. (Peng 2005) had previously shown Human PBMC to be an appropriate model to study the proinflammatory effect of CpG and non-CpG ODN.

PBMC (5×10^5^ cells/well) from 5 human volunteers were isolated and treated with either ISIS 104838 or ISIS 353512 within 2 hours of collection with increasing concentrations of ASOs from 0.4 μM to 100 μM. ISIS 104838 is another MOE ASO that had been through toxicology and showed a lesser degree of constitutional symptoms and ISRs(Sewell, Geary et al. 2002). The cells were incubated for 24 hours upon which time, the cell supernatant was collected and the levels for 17 cytokines and chemokines were measured using the MSD platform. Two distinct patterns emerged: IL-6, IL-10 and to a lesser extent TNF-α levels were clearly elevated in cells treated with ISIS 353512 relative to the other ASO tested (figure 2). While the magnitudes of the increase varied from donor to donor, the pattern remains consistent. IL-10 elevation was highest at the lowest dose tested (0.4 μM). IL-6 also showed a bell curve response to ISIS 353512, albeit less pronounced. In contrast, TNF-α displayed a more dose-responsive increase but with less separation between cell treated with ISIS 353512 and ISIS 104838. The other pattern of response to ASO treatment to emerge is a sequence-independent but ASO treatment dependent elevations in chemokines MDC or MIP-1β. Other cytokines and chemokines analyzed (see material and methods) showed no clear changes (data not shown). In follow up studies, the focus of the analysis centered around IL-6 and to a lesser extend IL-10 and TNF-α.

**Figure 2:**
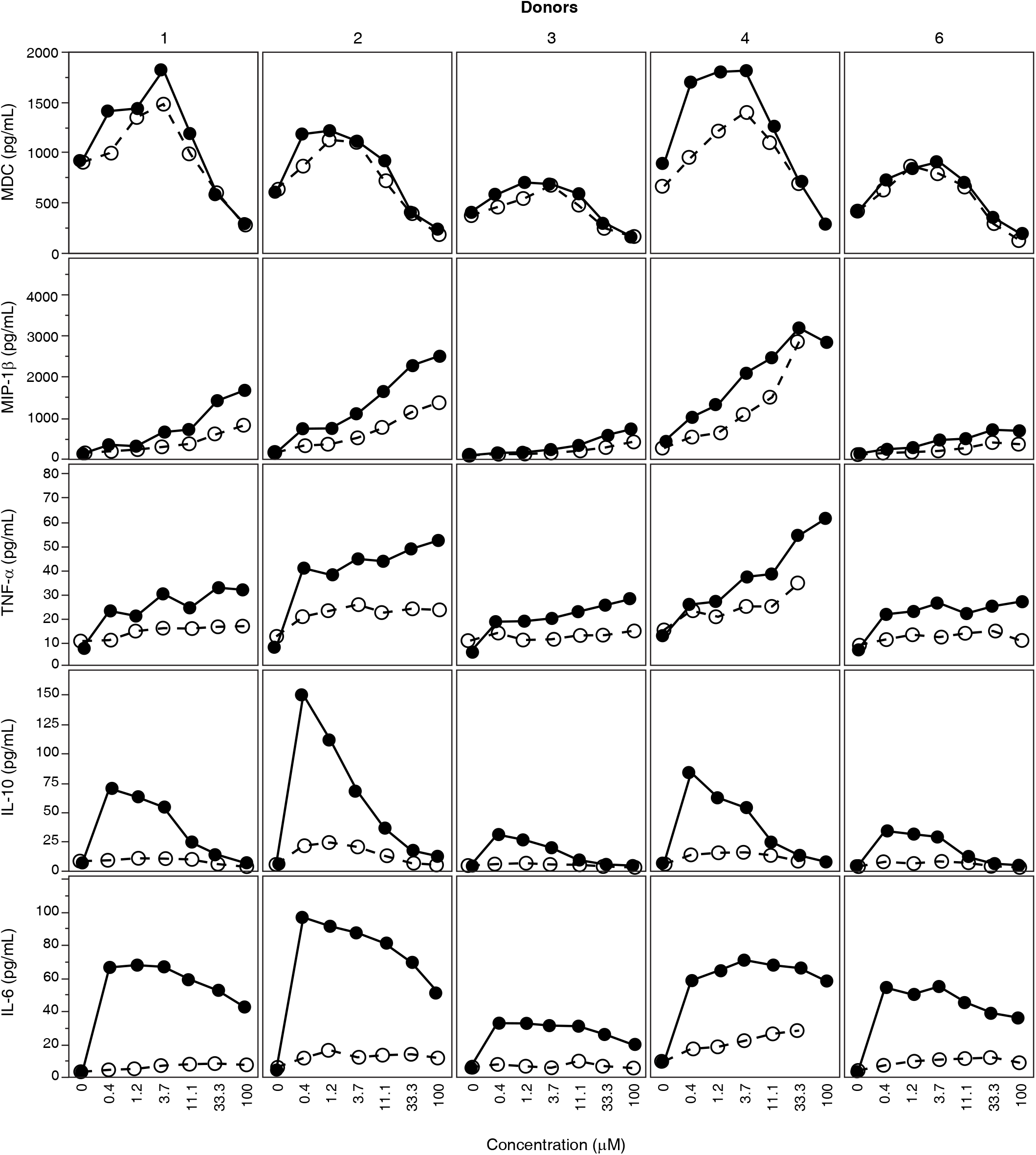
HsCRP level from a single dose of ISIS 353512 Phase I trial in healthy volunteers (a) Kinetic of serum human CRP upon iv dosing of ISIS 353512. ISIS 353512 (100mg) or placebo was administered on days 1. Dotted line represents the upper limit of normal at 8 mg/L (b) average of each dose group. (c) Serum IL-6 levels were measured by ELISA at prior to dosing and 8 hours after the first dose. n=3 per dose level

Next, we sought to investigate the range of responses of hPBMC exposed to either ISIS 353512 or ISIS 104838 of a broader set of healthy human blood donors (n=47). PBMC from healthy volunteers were isolated and plated cells at 5×10^6^ cells/mL and treated them with increasing concentrations of ASOs for 24 hours before harvesting the supernatant. Considering previous experiments conducted with a starting dose of 0.4 μM which indicated that IL-6 and IL-10 productions was already at its maximum at this dose level, a lower concentration range of MOE ASO from 0.019 μM to 80 μM was explored. The levels of IL-6, IL-10 and TNF-α were measured in the cell supernatant using MSD (supplemental figure 1). Overall, the hPBMC from various donors exhibited a very broad range of IL-6 production in response to ISIS 104838 and ISIS 353512 and fell into 2 broad categories: Discriminator and non-discriminator donors. Depending on the donor, the IL-6 response of hPBMC to ISIS 353512 peaks between 0.078 μM and 0.31 μM. To classify donors either as discriminators and non-discriminators, the ratio between the IL-6 level produced in response to 0.31 μM of ISIS 353512 and the IL-6 produced in response to 0.31 μM of ISIS 104838 were calculated (figure 3a). Donors with a ratio of 4 or greater were deemed to be discriminators (figure 3a). Based on this arbitrary threshold, hPBMC isolated from 21 discriminators donors could very easily separate ISIS 353512 from ISIS 104838, with ISIS 353512 consistently producing the highest increase in IL-6 (supplementary figure 1a and figure 3b). In contrast, hPMBC from other 26 donors (non-discriminator) failed to show much separation between the 2 MOE ASOs. When IL-6 levels were normalized for each donor to the untreated level, donors classified as discriminators can produce a much greater level of IL-6 in response to ISIS 353512 than ISIS 104838 relative to untreated IL-6 levels (figure 3c). In contrast, hPBMC from non-discriminator donors tended to have a higher baseline production of IL-6 (supplemental figure 3 and figure 3b) but much lower dynamic range of IL-6 production between ISIS 104838 and ISIS 353512 (figure 3c).

**Figure 3:**
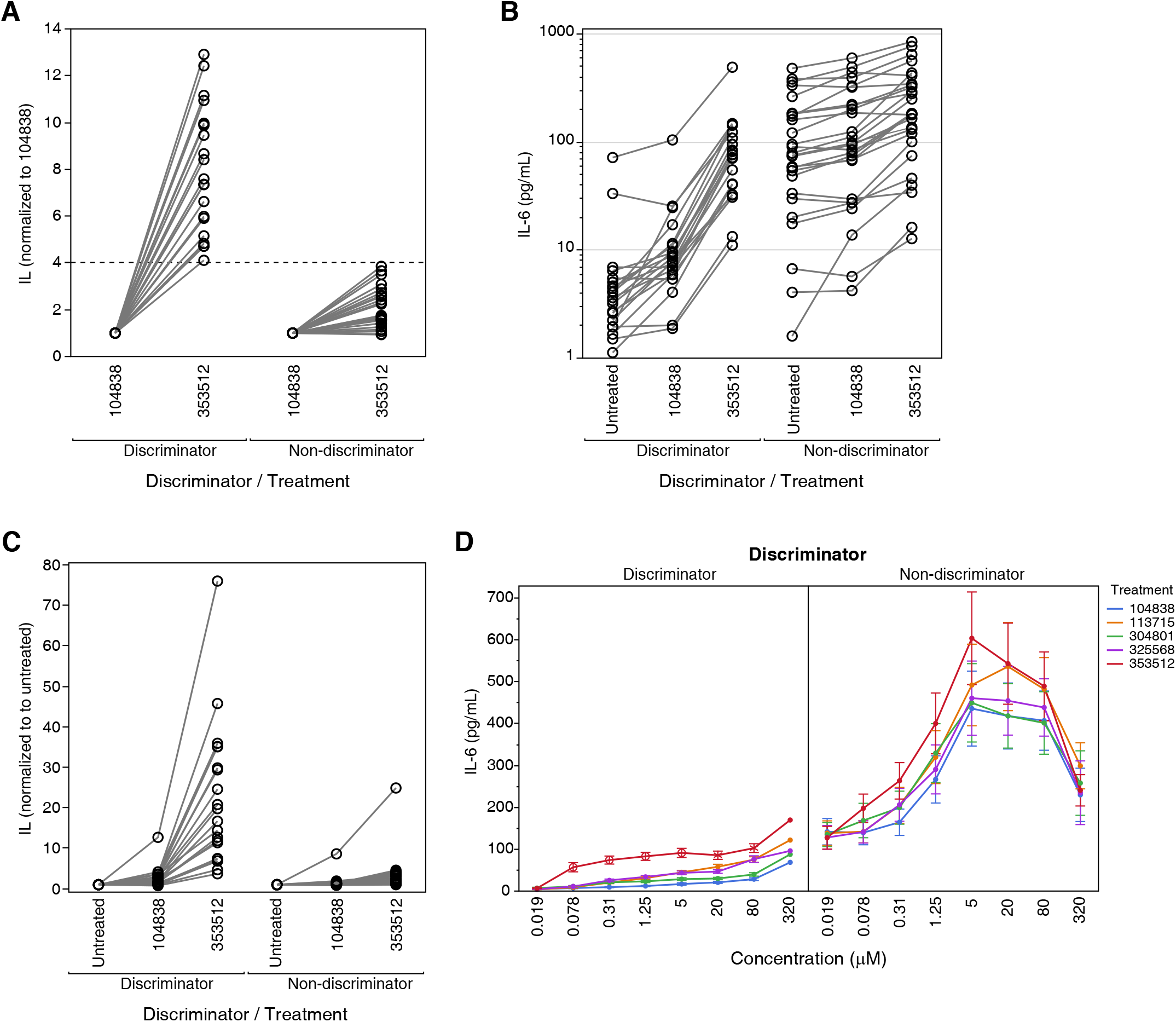
IL-6, IL-10, TNF-α, MDC and MIP-1β levels following treatment of hPBMC from 5 human volunteers treated with ISIS 104838 (open circles), ISIS 353512 (closed circles) for 24 hours. Cytokine and chemokine protein levels were measured using MSD.

To further validate hPBMC ability to identify inflammatory ASOs, we selected 3 additional MOE ASOs ASOs which had previously undergone phase I or phase II clinical trials (ISIS 104838 (NCT00048321)(Sewell, Geary et al. 2002), ISIS 113715 (NCT00330330), ISIS 304801 (Volanesorsen) (Graham, Lee et al. 2013) and ISIS 325568 (NCT00519727) (van Dongen, Geerts et al. 2015)) and found to have comparatively milder constitutional symptoms than those induced by ISIS 353512 (data not shown). Although hsCRP or IL-6 were not among the endpoints collected during the clinical trials with those ASOs, a retrospective review of constitutional symptoms and other safety related endpoint suggested that those MOE ASOs were better tolerated with respect to constitutional symptoms and ISR compared to ISIS 353512 (data not shown). In addition to ISIS 104838 and ISIS 353512, those ASOs were used to treat hPBMC cells from donors previously classified as discriminator and non-discriminator. While discriminator donors hPBMC treated with ISIS 353512 cells produced a significant increase in IL-6 (figure 3d) relative to ISIS 104838 at several of the ASO concentration tested, the other ASOs (ISIS 113725, ISIS 304801 and ISIS 325568) demonstrated marginal increase in IL-6 levels relative to ISIS 104838, none of which were significant. In contrast, ISIS 353512 (or ISIS 113715, ISIS 304801 and ISIS 325568) treatment of non-Discriminator donor hPBMC resulted in a similar degree of increase in IL-6 relative to ISIS 104838 at any of the ASO concentration tested. Interestingly, the level of IL-6 of in the cell culture supernatant from discriminator donor after 24 hours in absence of oligonucleotide treatment was markedly lower in most cases than non-discriminator donors (figure 3b, 3d and supplemental figure 3).

In contrast to IL-6, for which only treatment of hPBMC from discriminator donors enable to identify ISIS 353512 as distinctly proinflammatory relative to ISIS 104838, IL-10 levels were also significantly elevated at lower doses of ISIS 353512 in hPBMC from non-discriminator donors (supplemental figure 2). To a lesser extend ISIS 113715 and ISIS 325568 also caused some significant increase in IL-10 levels relative to ISIS 104838 but at doses higher than that of ISIS 353512.

### Reproducibility of donor specific response

Some hPBMC (discriminator) donors have been tested many times with ISIS 353512 and ISIS 104838. In figure 4, we compare the IL-6 production in response to ISIS 104838 or ISIS 353512 stimulation of hPBMC from 6 different donors in 5 independent runs of the assay. While the pattern of response remains fairly consistent over time for some donors (e.g., #52), most donors display a broad range of response to ISIS 35512. However, in all runs and all donors, stimulation with ISIS 353512 always lead to a greater production of IL-6 than stimulation with ISIS 104838.

**Figure 4:**
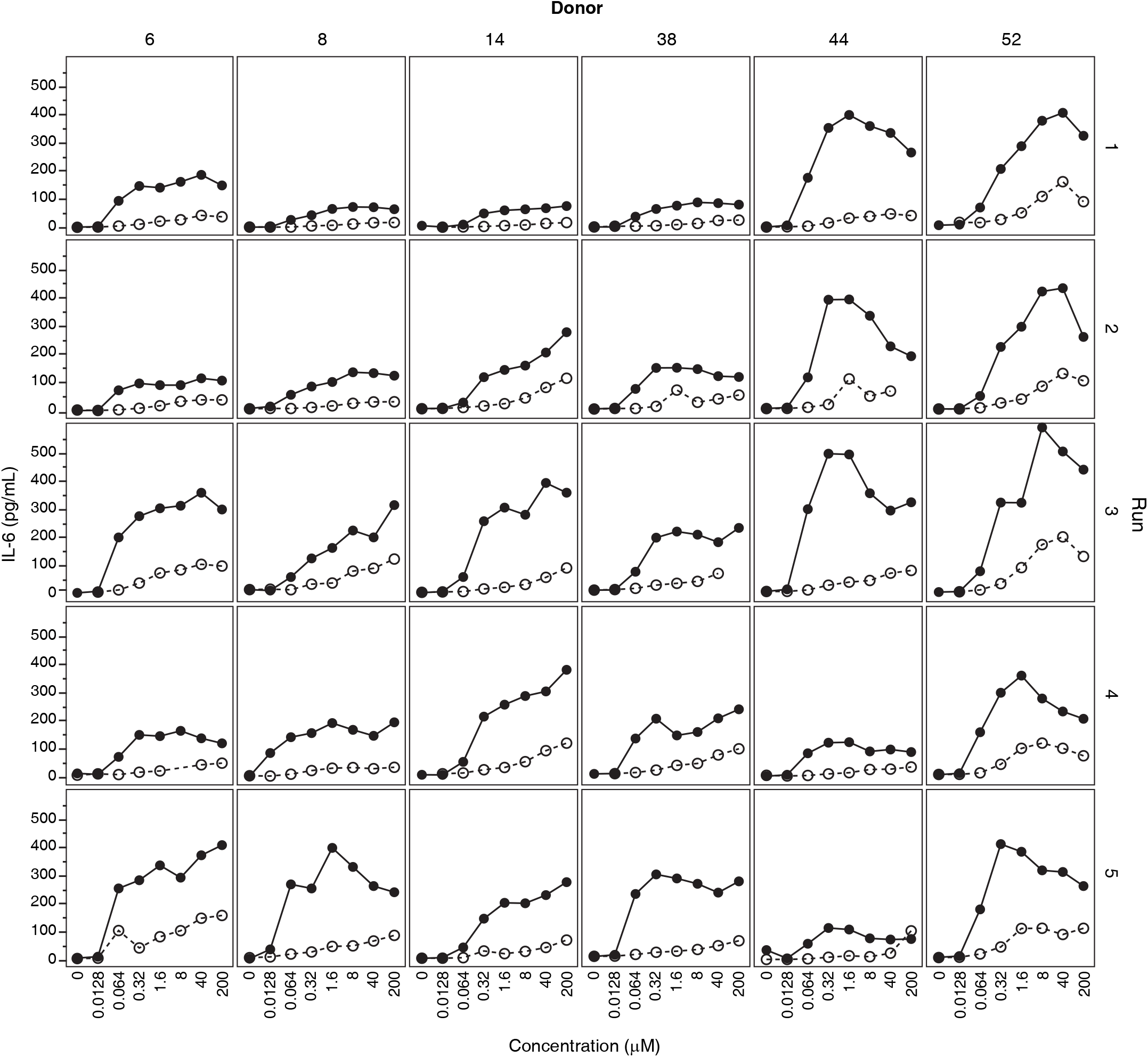
(a-c) Screening of hPBMC isolated from 47 healthy human volunteers. IL-6 level measured by MSD profiling following 24 hours incubations of hPBMC with 0.31 μM ASO treatments. 21 Donors were deemed capable of discriminating ISIS 353512 from ISIS 104838 response, while 26 Donors could not discriminate ISIS353512 from ISIS 104838 with confidence. Each line represents a different donor. (a) IL-6 level of ISIS 353512 at 0.31 μM normalized to ISIS 104838 at 0.31 μM (b) IL-6 levels for hPBMC dosed with 1.25 μM ASO (c) IL-6 level of ISIS 35512 and ISIS 104838 at 1.25 μM normalized to untreated level. IL-6 levels (d) measured concurrently by MSD multiplex profiling in the supernatant of hPBMC of 21 discriminator and 26 non-discriminator donors following 24 hours incubations of hPBMC with increasing concentration of 5 ASO treatments. Standard error of the mean is shown.

### PBMC isolated from Cynomolgus monkeys are an unsensitive to ISIS 353512 stimulation

As stated previously, preclinical safety studies conducted in mice or NHP did not identify the proinflammatory potential of ISIS 353512. Therefore, we sought to compare the effect of ISIS 353512 treatment in PBMC isolated from either human volunteers or naïve cynomolgus monkeys. NHP PBMC were isolated and exposed to ISIS 353512 or ISIS 104838 under condition very similar to those employed for the human PBMC assessment. After 24 hours exposure, neither of the two MOE ASOs assayed induced an increase in any of the cytokine or chemokine measured at ASOs concentrations overlapping those capable of eliciting an inflammatory response in hPBMC (using a panel shown to be cross reactive to NHP) (figure 5a). We sought to confirm that under the experimental conditions selected for the treatment, NHP PBMC retained their sensitivity to several well characterized proinflammatory stimuli. Responsiveness of monkey PBMCs to LPS, poly I:C and a CpG ODN (ODN 2395) (Roda, Parihar et al. 2005) was demonstrated thereby confirming that while functional in their ability to respond to the range of TLR agonist including TLR9 agonist (ODN 2395), NHP PBMC appeared to lack the ability to respond to more moderately proinflammatory MOE ASO such as ISIS 353512 in contrast to hPBMC (figure 5b). In addition, the magnitude of IL-6 production as measured using human anti-IL-6 detection confirm that the cross reactivity of the assay.

**Figure 5:**
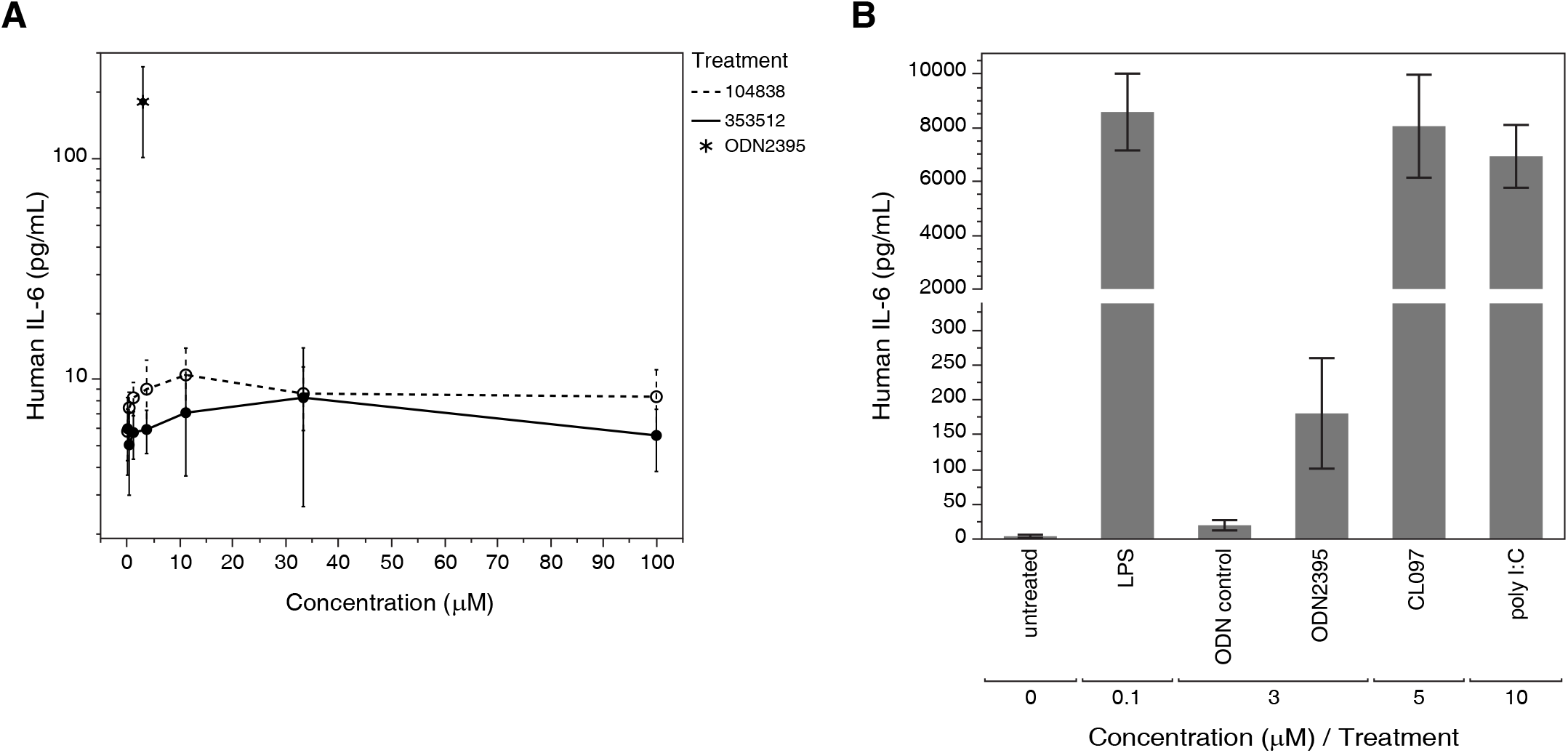
Reproducibility of IL-6 production in response to treatment to ISIS 104838 and ISIS 353512. PBMC from 6 discriminator donors were treated with either ISIS 104838 (open circles) or ISIS 353512 (closed circles) on 5 different occasions.

### B cells are producing IL-6 in response to ISIS 353512

Next, we sought to identify which cell type(s) among the hPBMC would be principally responsible for mediating the response to ISIS 353512. We had previously hypothesized that the most likely receptor mediating the inflammatory response would be TLR9 based on its recognition of CpG ODN (5).

In humans, two cell populations are mostly expressing TLR9: B cells and plasma dendritic cells. B-cells are mostly responsive to CpG type B ODN which have a structure most similar to 2’MOE ASOs in that they lack a PO central CpG-containing palindromic motif and a phosphorothioate-modified 3’ poly-G string present in CpG type A oligonucleotides to which plasma dendritic cells are more responsive (Krug, Rothenfusser et al. 2001, Vollmer, Weeratna et al. 2004). To ascertain the involvement of B cells, we used a mixture of magnetically labelled antibodies directed against all the main cell types present in hPBMC except for B-cells to negatively isolated B-cells from the rest of the hPBMC (figure 6). We also conducted the reverse experiment in which we also used magnetically labelled antibodies against B cells to depleted hPBMC from B-cells isolated from 2 different donors. We then proceeded to treat the 3 different populations (isolated and unlabeled B-cells, hPBMC depleted of B-cells or unmodified hPBMC) with either ISIS 353512 or ISIS 104838. Isolated B-cells retained their ability to respond to ISIS 353512 through the production of IL-6 (figure 6). IL-6 production of isolated B-cells in response to ISIS 104838 was somewhat increased relative to that of hPBMC treated with ISIS 104838 but nonetheless remained significantly lower than that to ISIS 353512. In contrast, B-cells treated with either ISIS 353512 or ISIS 104838 did not display much increased in IL-10 production relative to untreated cells in contrast to hPBMC treated with ISIS 353512 suggesting that the IL-10 production is a secondary event caused by the stimulation of other accessory cells (Couper, Blount et al. 2008). Further emphasizing the contribution of B-cells to the overall hPBMC response to inflammatory ASOs, hPBMC depleted of B-cells largely failed to show any increase in any of the cytokines assayed (figure 6).

**Figure 6:**
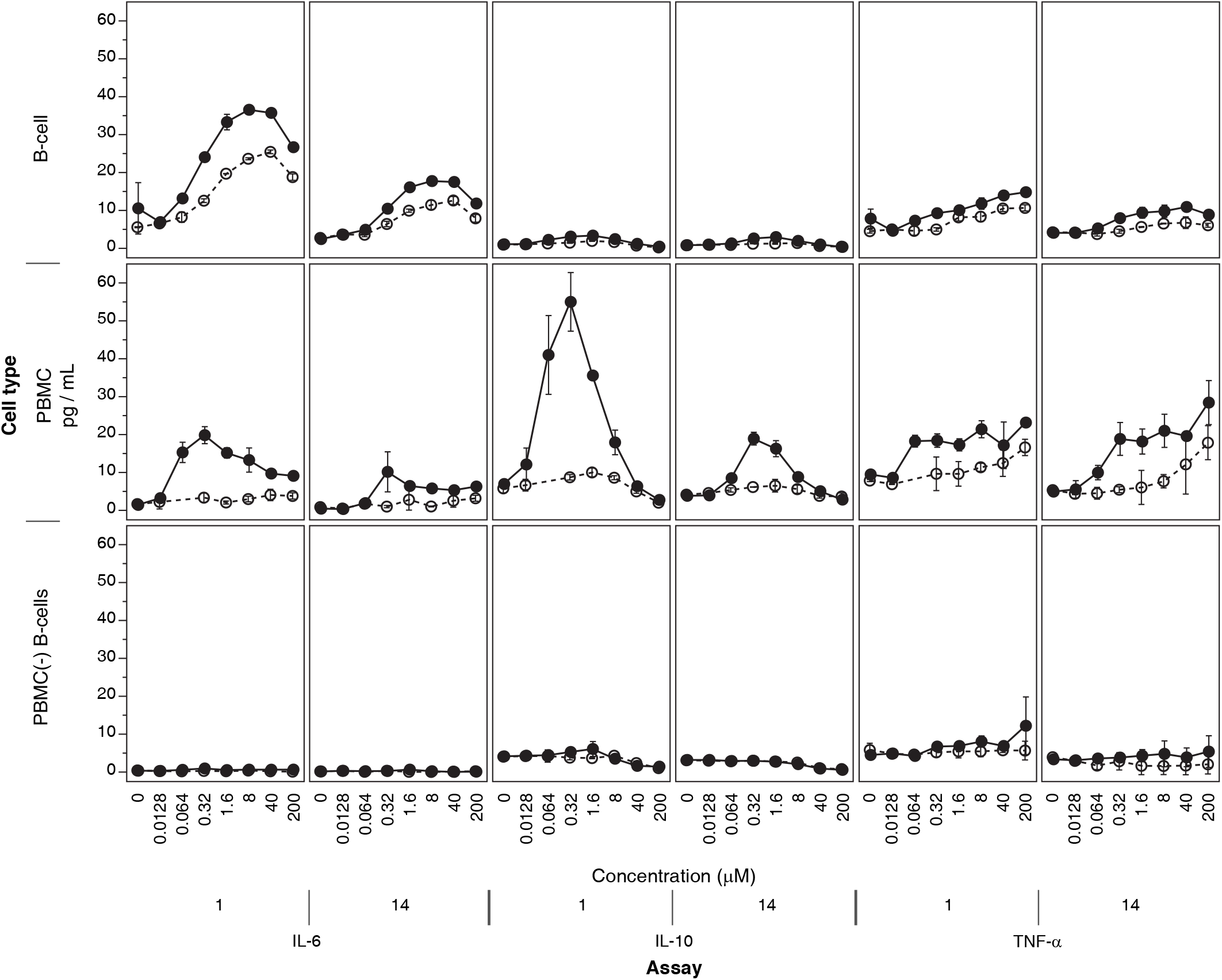
(a) PBMC isolated from cynomolgus monkeys were treated with either 3 μM ODN2395 (star) or increasing concentration of ISIS 353512 (closed circles) or ISIS 104838 (open circles). (b) PBMC isolated from cynomolgus monkeys were treated with several positive controls. IL-6 level measured by MSD multiplex profiling is shown (more analytes shown to be cross reactive were measured). Treatment were conducted in triplicate. Error bars are representing the standard deviation of the mean.

### B-cell response to ISIS 353512 and ODN2006 is inhibited by chloroquine pretreatment

TLR9 signaling by proinflammatory CpG containing ODNs requires internalization of TLR9/ODN complex (Hong, Jiang et al. 2004, Leifer, Kennedy et al. 2004, Kuznik, Bencina et al. 2011). Once internalized, TLR9 cleavage and activation by cathepsin which requires endosomal acidification takes place. Suppression of endosomal TLR9 activation can be induced by inhibition of endosomal acidification using chloroquine, a weak base which accumulates in acidic compartments (Rutz, Metzger et al. 2004). This requirement for internalization and endosomal acidification is shared by TLR3, TLR7 and TLR8 which also share with TLR9 an ability to respond to nucleic acid-based pathogens-associated molecular patterns (PAMP) in contrast to other TLRs (TLR1, TLR2, TLR4, TLR5, and TLR6) which do not recognize non nucleic acid based PAMP (such as LPS (TLR4) and flagellin (TLR5)). In order to determine whether B-cells were responding to ISIS 353512 stimulation through TLR9 activation and signaling, hPBMC were pretreated with 10 μM Chloroquine which resulted in the complete loss of the IL-6 production by either ISIS 353512, ISIS 518477 (Paz, Hsiao et al. 2017) and ISIS 120704 (CpG, ODN 2006) (figure 7a). ISIS 120704 is CpG containing phosphorothioate ODN, while ISIS 518477 a 2’MOE gapmer ASO which does not contain any canonical CpG motif, have been previously shown to activate TLR9 signaling in rodents (Paz, Hsiao et al. 2017). hPBMC response to LPS (TLR4), Flagellin (TLR5) and other synthetic activators of non-nucleic acid PAMP responsive TLRs which does not depend on endosomal acidification as a prerequisite for activation is essentially unaffected. In contrast, TLR3, TLR7 and TLR8 activation by synthetic agonists was abrogated in presence of chloroquine (supplemental figure 4).

**Figure 7:**
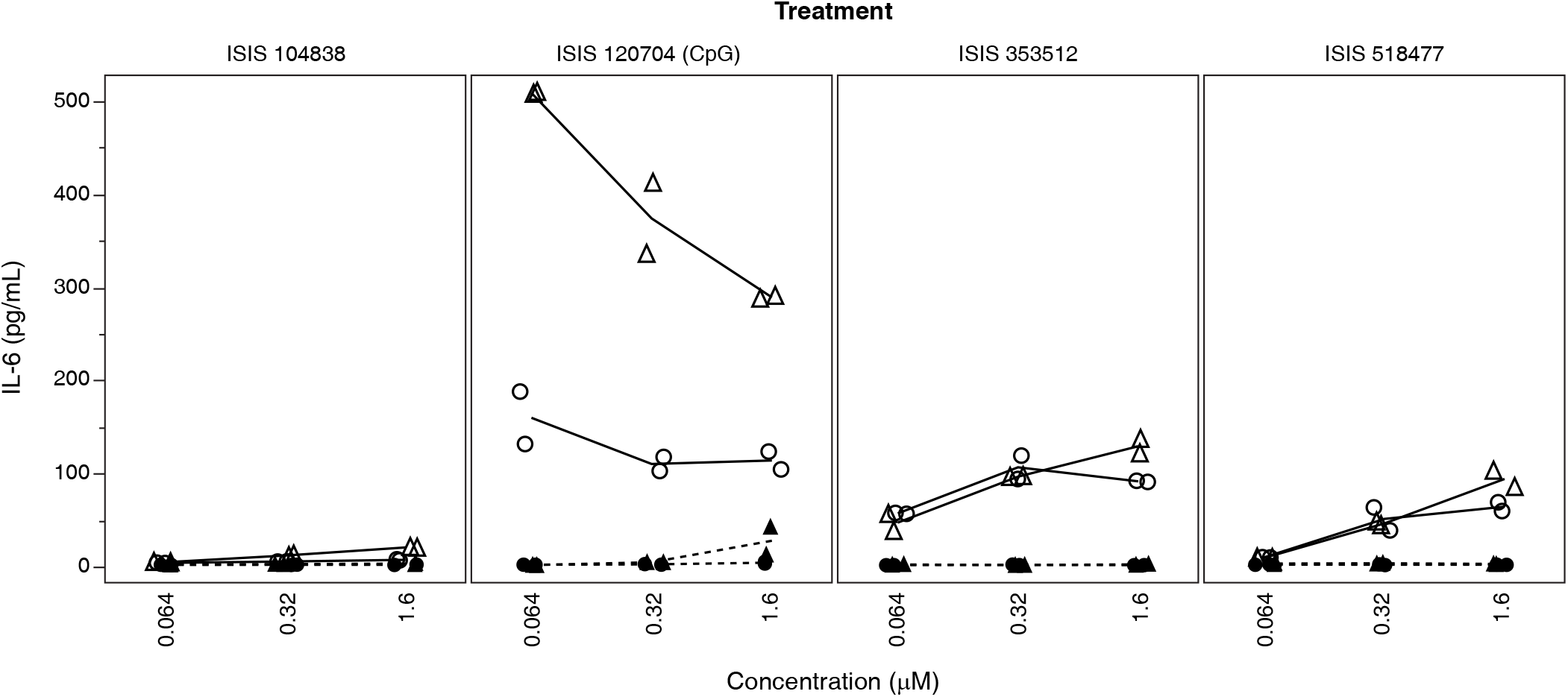
(top) B cells (isolated from PBMC – negative selection from donors 1 and 14) response to ISIS 353512 (closed circles) and ISIS 104838 (open circles) was compared to (middle) hPBMC or (bottom) hPBMC depleted of B cells (positive selection of B-cells. IL-6, IL-10 and TNFα was measured by MSD in the supernatant of the three cell populations following 24h exposed to ISIS 3535312 for 24 hours.

### ISIS 353512 immunostimulation depends on the presence of unmodified bases

ISIS 353512 uses a ‘gapmer’ design in which the sugars for 3 bases on the 3’ end and 3 bases on the 5’ end of the oligonucleotides are contains 2’MOE modifications. The remaining bases are nature DNA sugars. We hypothesized that in part the increase proinflammatory signal was mediated by the size of the unmodified ‘gap’. A major improvement in the pharmacological and tolerability profile of oligonucleotides was the addition of modified bases on the 5’ and 3’ end of the ODN. This resulted in increased stability, increased activity as well as decreased proinflammatory properties relative to fully unmodified phosphorothioate ODNs (Monteith, Henry et al. 1997). Indeed, the increased number of modified 2’MOE bases and therefore the reduction in gap size from 14 bases (ISIS 353512) to 10 bases (ISIS 330012) resulted in a profound reduction in IL-6 production by hPBMC from 2 donors (supplemental figure 6).

## Discussion

During the pharmaceutical development process, establishing the safety or tolerability of a new pharmacological entity in patients is paramount. This relies on establishing a therapeutic index or margin: Chiefly the ratio between the dose which would result in an unacceptable incidence of adverse events (whatever they maybe) and the dose which achieves the desired pharmacological effect designed to decrease the disease burden for the patient. One challenge in the nonclinical safety assessment is the translational relationship, or predictiveness of animals to humans. In most cases with ASOs transitioning into clinical trials, this approach has been reliable means of conducting the safety assessment. ISIS 353512 was an exception as it had been shown to be well tolerated in both rodents and NHP 3 months tolerability studies (figure 1), yet during the phase I/II human trials (NCT00734240), it became clear that the incidence of mild to moderated constitutional symptoms, along with increases in IL-6 and CRP at dose level required to demonstrate therapeutic efficacy were unacceptably high. While the clinical findings were transient and did not result in severe injuries to the human volunteers, the discrepancy between the outcome of animal studies and those clinical findings demonstrated the necessity to include a human based measure of inflammation to the nonclinical safety assessment process of ASOs. This is now a routine part of the compound selection and safety testing at Ionis Pharmaceuticals.

Past experience has shown mice to be a sensitive species suitable to screen out nonspecific and proinflammatory effects of non-CpG ASO prior to the initiation of human clinical trials (Monteith, Henry et al. 1997). This approach resulted in multiple ASOs translating successfully into the clinic with acceptable proinflammatory profiles. In contrast, ISIS 353512 phase I findings revealed exaggerated proinflammatory phenotypes (volunteers treated with ISIS 353512 showed a higher incidence in mild to moderate constitutional symptoms such as fever, chills, asthenia, feeling hot and feeling cold,… as well as Injection site reactions including injection site pain, erythema, itching and swelling, elevated IL-6 and CRP observed shortly after the first dose of ISIS 353512 relative to placebo treated volunteers). Those findings were not predicted by either mice or monkey studies in which evidence of a proinflammatory response was minimal and similar to previous ASOs which successfully entered clinical trials. The nature of the clinical findings suggested a B cell mediated response might be at play while the nature of the stimuli which is synthetic DNA-like suggested an innate immune receptor, likely TLR9, might be responsible for the proinflammatory response observed in the study. We and other authors had previously demonstrated that such a receptor, TLR9, mediated the inflammatory response to some ASOs in mice despite the absence of any kind of CpG motif in those ASOs (Vollmer, Janosch et al. 2002, Vollmer, Weeratna et al. 2004, Paz, Hsiao et al. 2017).

Oligonucleotides containing CpG motifs are known to elicit robust proinflammatory responses from rodent to humans (Krieg, Yi et al. 1995, Krieg 1999, Bauer, Kirschning et al. 2001). Furthermore, TLR9-mediated DNA recognition and cell distribution has been shown to vary between species. In humans, TLR9 is predominantly expressed in B cells and plasma dendritic cells, while in rodents, TLR9 is expressed more widely including myeloid immune cells. The optimal CpG motifs capable of activating TLR9 have also shown to vary slightly from one species to the next (Pohar, Kuznik Krajnik et al. 2015, Pohar, Lainscek et al. 2015). Therefore, we hypothesized that despite the absence of canonical CpG motif, ISIS 353512 might be inducing a TLR9 mediated response involving primarily B cells (Vollmer, Janosch et al. 2002, Elias, Flo et al. 2003, Vollmer, Weeratna et al. 2004) and that the lack of ISIS 353512 proinflammatory stimulation in rodent may stem from species-specific recognition of CpG and non-CpG ODN (Campbell, Cho et al. 2009).

As with CpG oligonucleotides exemplified by ODN2006 (ISIS 120704), treatment of PBMC collected from 5 human volunteers with ISIS 353512 resulted in IL-6 increase 24 hours after the initiation of the treatment in the majority of the PBMC sample tested (figure 3, figure 7). In contrast, this elevation in IL-6 was either minimal or not observed following exposure with ISIS 104838. The clinical experience with ISIS 104838 suggested that it was overall better tolerated in human volunteers than ISIS 353512 with a lower incidence or severity of flu like symptoms (Sewell, Geary et al. 2002). As ISIS 353512, ISIS 104838 was found to be well tolerated in rodent and non-human primate studies. As such, ISIS 104838 was selected as being representative of oligonucleotides with an acceptable tolerability profile in human volunteers. In parallel to testing those oligonucleotides, in hPBMC, we also tested them using whole blood collected from the same volunteers. While treating whole blood samples with those oligonucleotides also resulted in differential increase in IL-6 which clearly discriminated ISIS 104838 from ISIS 353512 (data not shown), we settled on optimizing a tolerability screening assay using hPBMC due to the ability to standardize the starting cell number to achieve greater consistency of response.

In follow up investigations, we sought to investigate the consistency of the response to ISIS 353512 across a larger number of hPBMC samples. Surprisingly, the response of the hPBMC to ISIS 353512 could be assigned to 2 categories (figure 3 and supplemental figure 1, 2 and 3): 21 donor samples demonstrated a good separation between ISIS 104838 and ISIS 353512 at a dose of 0.31 μM with a ratio of IL-6 levels of 4 or more between the 2 oligonucleotides (these donors were referred to as discriminators). In contrast, 26 donors were largely unable to discriminate those oligonucleotides (with an ISIS 353512/ISIS 104838 IL-6 ratio below 4) (these donors were referred to as non-discriminators). While the threshold is arbitrary, it is supported by the level of IL-6 production in absence of treatment. Nineteen out of 21 discriminator donors had IL-6 level between 1 and 10 pg/mL after 24 hours of incubation without oligonucleotides (supplemental figure 3). In contrast, under the same conditions, non-discriminator donors exhibited IL-6 greater than 20 pg/mL in 23 out 26 cases. The basis for the differential sensitivity of PBMC from human volunteers to ISIS 353512 remains unclear and does not seem to translate to the clinical experience (albeit limited in scope) in which all volunteers treated with ISIS 353512 exhibited some increase in hCRP. For ethical reasons, we were not able to ascertain whether the discrimination could be translated in vivo in a clinical setting. We were unable to test retrospectively the responsiveness of hPBMC from volunteers enrolled in the clinical trials to further explore the connection between in vitro and in vivo response. Regardless, the presence of 2 types of in vitro responses to ISIS 353512 highlighted the need for prescreening donors to identify samples with the optimal response profile to ISIS 353512 before screening novel oligonucleotides. For donors classified as discriminators, while ISIS 353512 consistently led to a greater increase in IL-6 relative to ISIS 104838, the absolute level of IL-6 productions varied markedly from one run to the next (figure 4) which illustrated the need to include some reference compounds such as ISIS 104838 and ISIS 353512 to be able to determine whether the PBMC from a given donor are behaving as required.

Typically, screening using hPBMC to determine the likelihood of novel compounds to produce an adverse proinflammatory response is performed at least two different (discriminator) donors. Interpreting the screening data to determine whether a novel compound should move forward largely relies on the relative ranking of compounds rather than absolute pass or fail threshold. Any oligonucleotide which elicits an IL-6 which nears or exceeds that of ISIS 353512 in at least 1 donor is typically eliminated. Similarly, any oligonucleotide which produce an increase in IL-6 that nears or is below that of ISIS 104838 in both donors is deemed to have a very low risk of producing an adverse inflammatory response in patients. Typically, any oligonucleotide which exhibit a relative increase in IL-6 halfway between ISIS 104838 and ISIS 353512 is treated with caution.

As such, A follow-on MOE ASO to ISIS 353512, ISIS 329993, also designed to reduce levels of hsCRP mRNA (and protein) was administered to human volunteers in both single and multiple ascending dose double-blind randomized phase I clinical trial at a starting dose of 50 mg (Noveck, Stroes et al. 2014). ISIS 329993 is targeting a different region of the CRP mRNA transcript and therefore has distinct sequence compared to ISIS 353512 and also has a 5-10-5 MOE Gapmer configuration (supplemental table 1). In contrast to ISIS 353512, treatment with ISIS 329993 did not cause an increase in hsCRP in human volunteers at dose up to 400 mg/week SC relative to placebo treated volunteers. In addition, clinical observations of the volunteers treated with ISIS 329993 showed a marked reduction in incidence and severity of constitution symptoms such as fever, chills, aches,… relative to ISIS 353512. ISIS 329993 was well tolerated at doses in excess of those dose tested with ISIS 353512 and progressed to phase II (NCT01710852 and NCT1414101). ISIS 329993 caused little to no increase in CRP level even at dose of 600 mg. The minimal immune stimulation seen in the clinical trial was paralleled by the minimal induction in IL-6 production by hPBMC treated with ISIS 329993. Besides ISIS 353512 and ISIS 329993 having different sequences, ISIS has 14 unmodified bases with phosphorothioate linkages against 10 for ISIS 329993 or ISIS 104838. When the number of unmodified bases was reduced from 14 (ISIS 353512) to 10 (ISIS 330012), IL-6 production by hPBMC treated with ISIS 330012 was very similar to that of hPBMC cells treated with ISIS 104838 (supplemental figure 6) indicating that the number of unmodified bases with phosphorothioate linkages is an important contributor the immune stimulation by ASOs. In recent years, most ASOs that have transitioned to a preclinical tolerability study in NHP and eventually to a clinical study have a ‘gap’ of 10 unmodified bases or less with ‘wings’ with modified bases that can vary in nature (mostly 2’MOE or cEt, (Seth, Siwkowski et al. 2009)). As a result, 130 5-10-5 MOE gapmer ASOs scheduled to be tested in NHP preclinical studies have been tested in the hPBMC assay prior to the start of those studies. IL-6 profiling from hPBMC treated with those ASOs indicated that even with a constant number of unmodified bases, ASOs with difference sequence composition but identical chemistry design can display a wide range of IL-6 induction, in some cases exceeding that of ISIS 353512 by more than one order of magnitude despite the absence of canonical CpG motif in the sequence indicating the both the number of unmodified bases and the nature of the sequences are two fundamental modulators of the potential to trigger IL-6 stimulation (supplemental figure 7).

We sought to gain a better understanding of the mechanism underpinning the production of IL-6 following treatment with ISIS 353512. As mentioned previously, we hypothesized that the mechanism might be in part overlapping with that of canonical CpG oligonucleotides. Both B-cells and dendritic cells are expressing TLR9. Depending on the nature of the CpG motif, oligonucleotides, the response will be either mostly driven by B-cells or dendritic cells (Krug, Rothenfusser et al. 2001, Vollmer, Weeratna et al. 2004). Three types of stimulatory CpG ODNs have been characterized, type A, B and C, which differ in their immune-stimulatory activities (Krug, Rothenfusser et al. 2001, Marshall, Fearon et al. 2005). ODN2006 (ISIS 120704) is a 24-mer ODN with a full phosphorothioate backbone and belongs to the type B CpG ODNs which contain one or more unmethylated CpG dinucleotides in particular sequence contexts (CpG motifs) and are recognized by human TLR9. Type B CpG ODNs strongly activate B cells but stimulate weakly IFN-α secretion. Type A CpG ODNs are characterized by a phosphodiester central CpG-containing palindromic motif and a phosphorothioate 3’ poly-G string. They induce high IFN-α production from plasmacytoid dendritic cells. Type C CpG ODNs combine features of both types A and B. In the present case, the structure of ISIS 353512 loosely resembles CpG type B such as ISIS 120704 (Hartmann, Weeratna et al. 2000, Bauer, Kirschning et al. 2001) in that ISIS 353512 does not include a 3’ poly-G string characteristic of type A and C CpG ODNs. The ability of ISIS 353512 to induce an inflammatory response was compared to ISIS 120704 in 9 different hPBMC donors (supplemental figure 8 a, b). ISIS 120704 induced a maximum 189 folds increase in IL-6 production at 0.064 μM relative to untreated hPBMC. In contrast, treatment with ISIS 353512 resulted maximum 37 folds increase in IL-6 production at 1.6 μM relative to untreated hPBMC. Finally, treatment with ISIS 104838 induced a maximum 11 folds increase in IL-6 production at 200 μM relative to untreated hPBMC. Despite the inadequate safety profile of ISIS 353512, the potency of ISIS 353512 to induce an inflammatory response is much lower than a CpG oligonucleotide such as ISIS 120704.

Next, we sought to examine whether the cellular response to ISIS 353512 might be driven by B cells. The ability of human PBMC and B cells in particular to mount an IL-6 response to CpG oligonucleotides treatment has previously been demonstrated (Cooper, Ahluwalia et al. 2008). Indeed, depleting PBMC of B-cells resulted in the complete abrogation of IL-6, IL-10 and TNF-α response in hPBMC from 2 separate donors (figure 6). When B cells were isolated and treated with ISIS 353512, the ability to produce an IL-6 response to the treatment was retained. Interestingly, the IL-10 production and to a lesser extend TNF-α remained absent suggesting that the IL-10 response observed in hPBMC is a secondary event caused by the stimulation of other accessory cells indicating an interaction between distinct cell populations in the hPBMC (Couper, Blount et al. 2008).

While treating hPBMC with ISIS 353512 consistently produces more IL-6 than ISIS 104838 in all the donors tested, this is achieved under specific collection conditions. Typically, PBMC are isolated from the blood of healthy volunteers and treated within a 3-hour window. Attempt to streamline the assay by either resting PBMC at 4°C overnight or freezing PBMC obtained from buffy coats resulted in a systematic loss of IL-6 following stimulation with ISIS 353512 (data not shown). This result was in contrast with the stimulation with a canonical CpG oligonucleotide such as ISIS 120704 for which resting or freezing PBMC prior the assay resulted in minimal loss of IL-6 induction (data not shown). The reason for this anergy could not be established. Despite the rapid anergy limiting our ability to fully dissect the mechanism, we gain some limited insight to the IL-6 production by ISIS 353512 by pretreating hPBMC with chloroquine. Chloroquine prevents the endosomal acidification and the activating cleavage of TLR7, 8 and 9 (Hong, Jiang et al. 2004, Latz, Schoenemeyer et al. 2004, Rutz, Metzger et al. 2004). Chloroquine treatment suppressed TLR7, TLR8 and TLR9 mediated inflammatory response by their respective canonical agonist (Imiquimod, ssRNA-40 and ISIS 120704 respectively). The same was also true for ISIS 353512 and ISIS 518477 where chloroquine treatment of hPBMC also substantially reduced the IL-6 production (figure 8). While we were unable to directly demonstrate that ISIS 353512 treatment leads to an inflammatory response that mediated through an interaction with TLR9, the circumstantial evidence (B-cells responsiveness and mitigation of the inflammatory response by Chloroquine) presented above strongly implicate a subset of the TLRs and TLR9 in particular as the key mediator.

In conclusion, we describe a simple in vitro screening paradigm to supplement rodent and nonhuman primate toxicology studies which enable the investigators to reliably identify oligonucleotides capable of eliciting an adverse inflammatory response specifically in humans prior the initiation of clinical trials thereby minimizing the risk to patients. This screening paradigm has a limited throughput due to its requirement for freshly collected human PBMC. In addition, the donor-to-donor variability reduces somewhat the robustness of the assay. Nonetheless, the nature of the cell responsive to ASO stimulation (B cells) and the likely receptor (TLR9) opens up a path forward to identify a cell line-based alternative which would replicate most of the feature of the hPBMC assay while increasing the throughput and consistency of the assay thereby enabling users to implement the cell line assay at a much higher throughput.

## Acknowledgement

The authors thank the donors who participated in this study; Erin Morgan for her help in critical review of an earlier version of the article; and Wanda Sullivan for assistance with formatting the figures and Angela Colabucci in proofreading of the article.

## Author Disclosure Statement

The authors are either current or former employees of Ionis Pharmaceuticals

---

**Figure 8**: Responses of hPBMC from 2 different human volunteers (44 (circle) and 52 (triangle)) to synthetic TLR agonists (a) or proinflammatory ODN stimulation (b) was tested with in absence (solid line and open symbols) or presence (dotted line and close symbols) of 10 μM Chloroquine.

