## Supplemental Figures 1-8 for "Early-Stage Identification and Avoidance of Antisense Oligonucleotides Causing Species-Specific Inflammatory Responses in Human Volunteers Peripheral Blood Mononuclear Cells"

Supplemental Figure 1

A

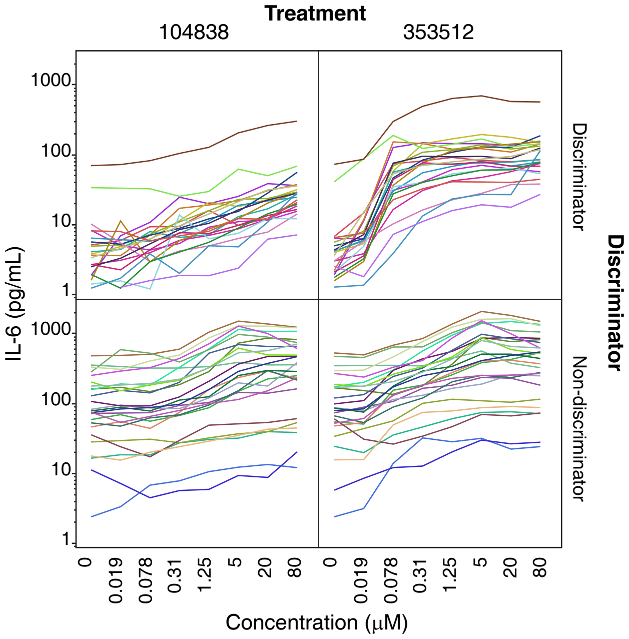


B

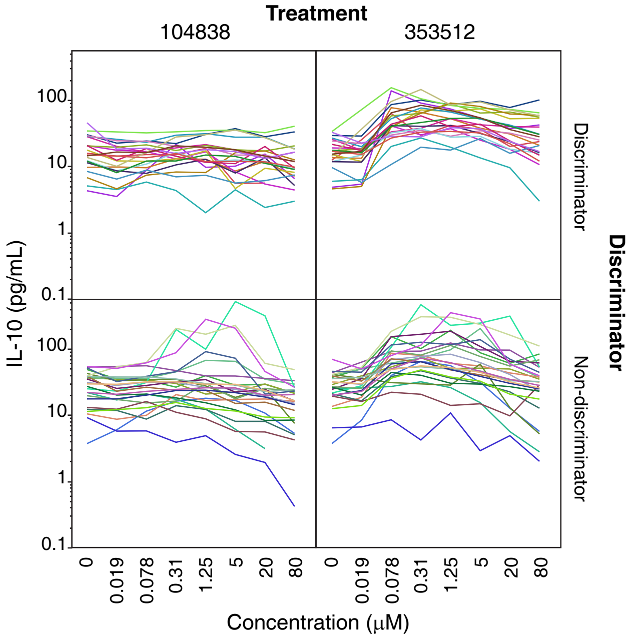


C

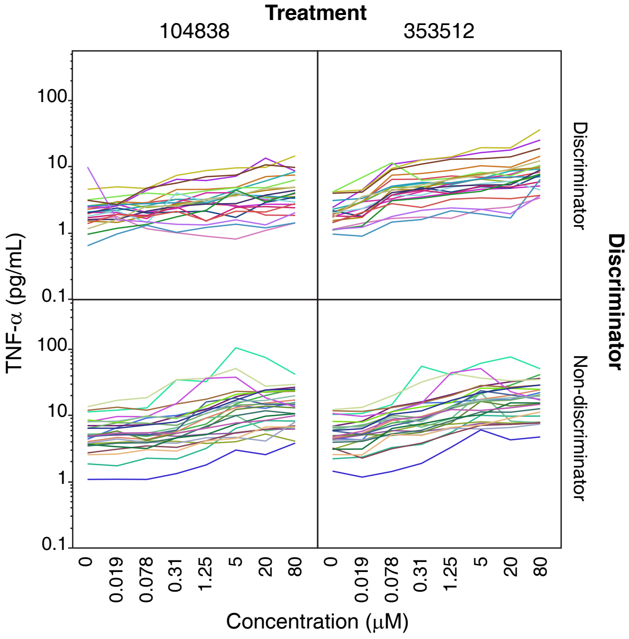

Supplemental Figure 2

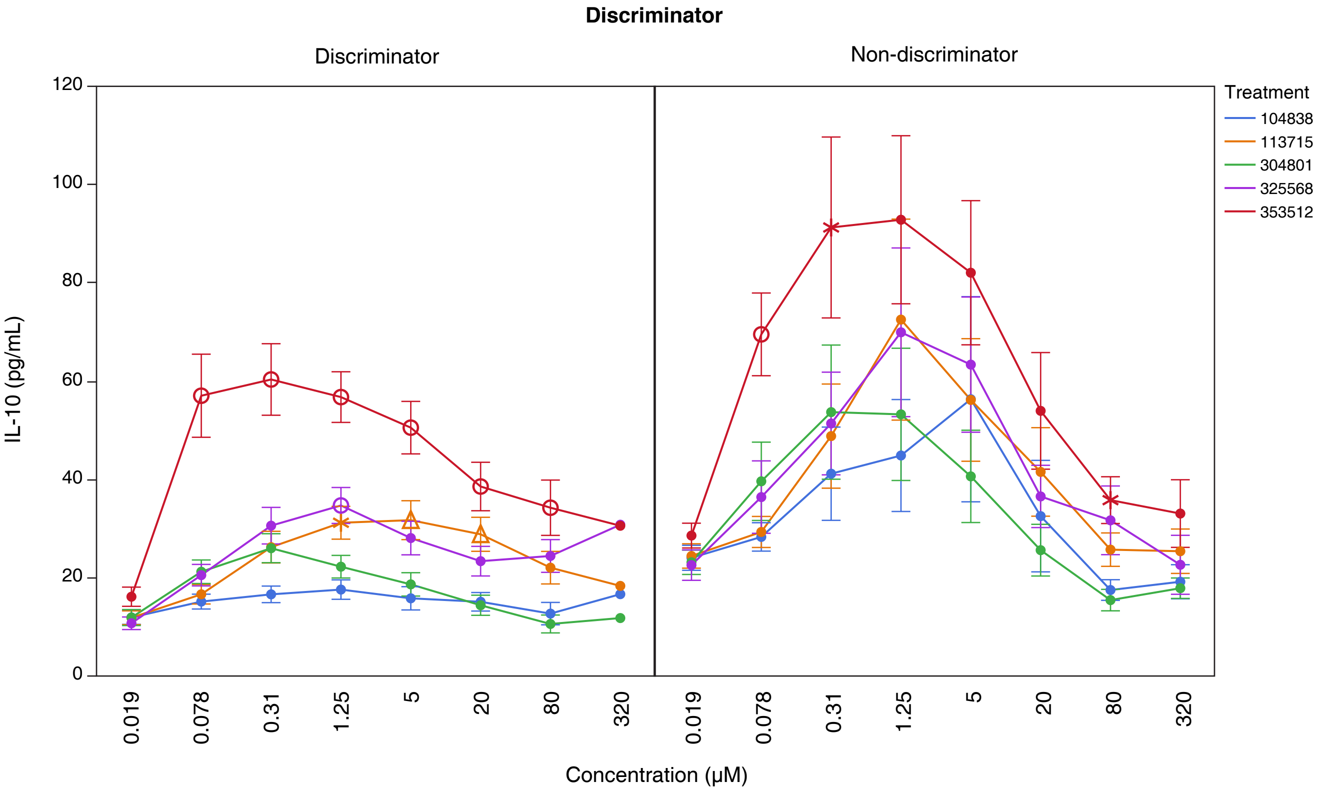

Supplemental Figure 3

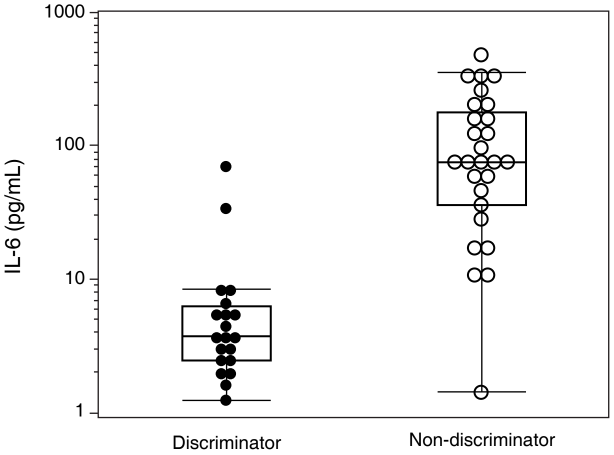

Supplemental Figure 4

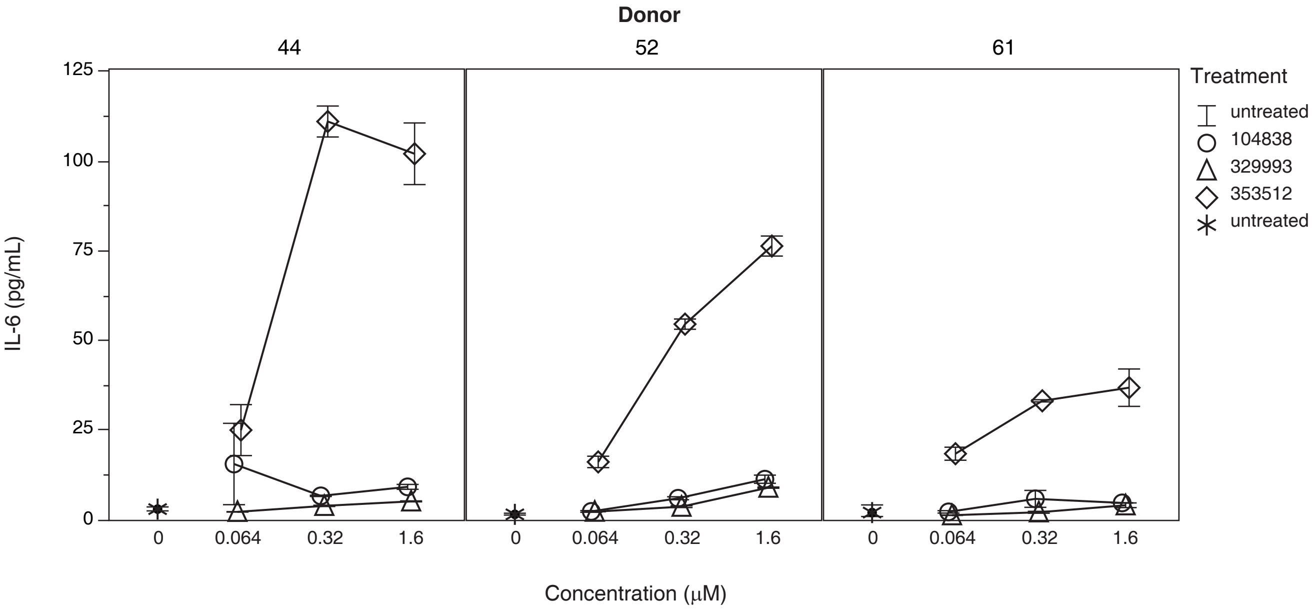

Supplemental Figure 5

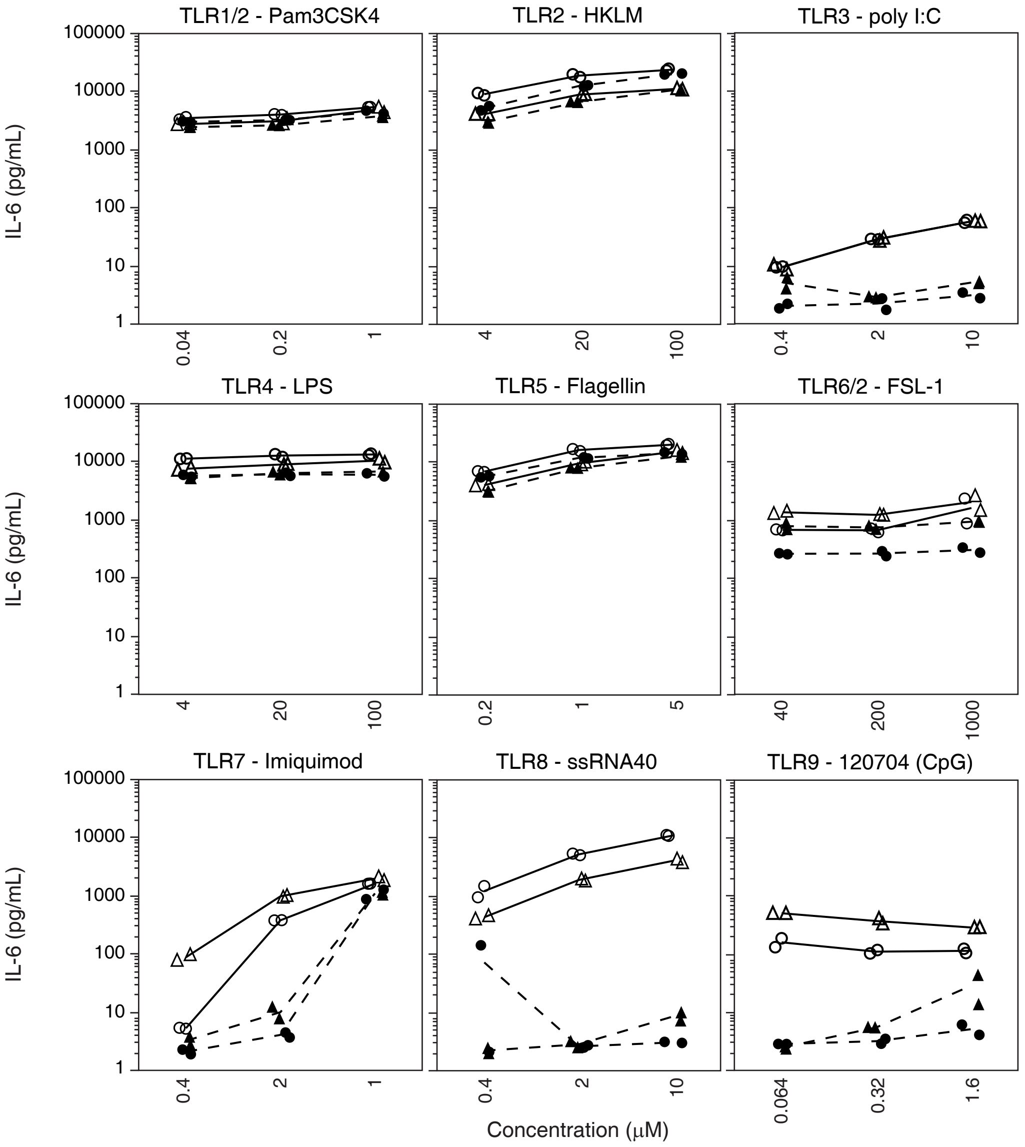

Supplemental Figure 6

**A**

| ODN ID | Full Sequence |
|---|---|
| ISIS 330012 | TsCsCsCsAsTsTsTsCsAsGsGsAsGsAsCsCsTsGsG |
| ISIS 353512 | TsCsCsCsAsTsTsTsCsAsGsGsAsGsAsCsCsTsGsG |

**B**

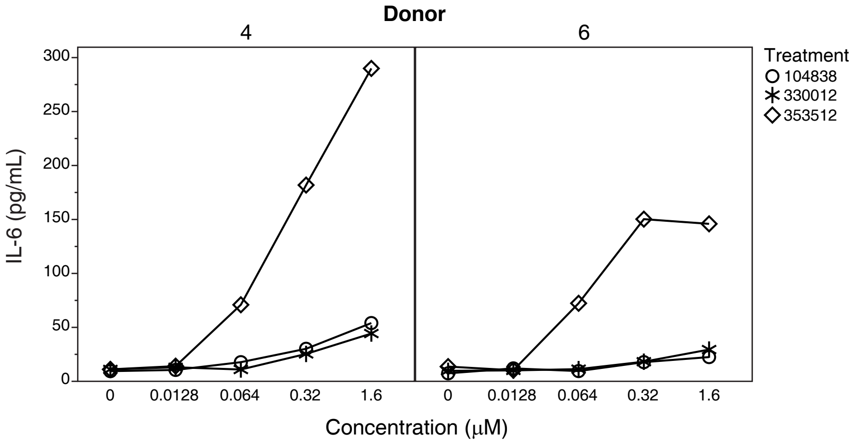

### Supplemental Figure 7

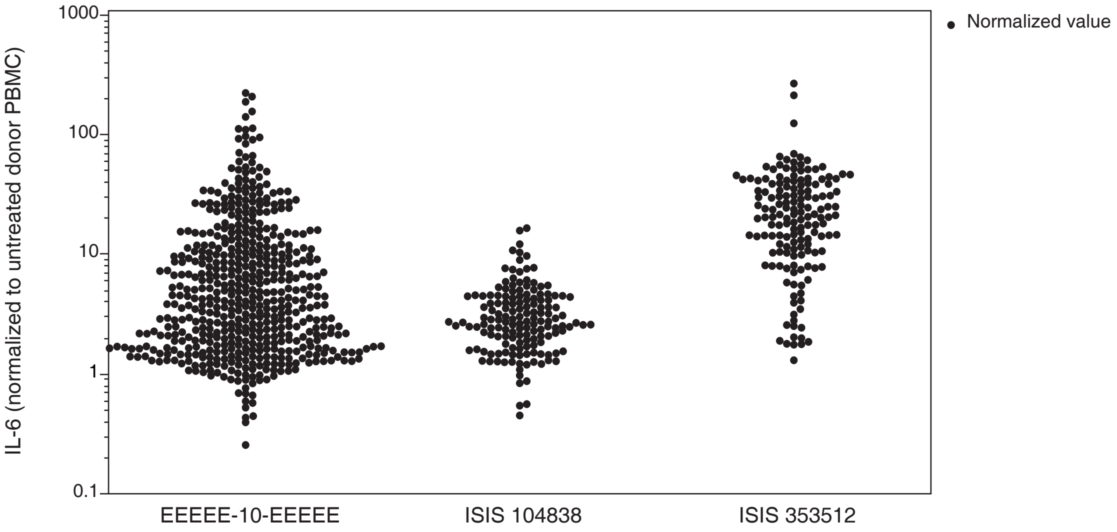

Supplemental Figure 8

**A**

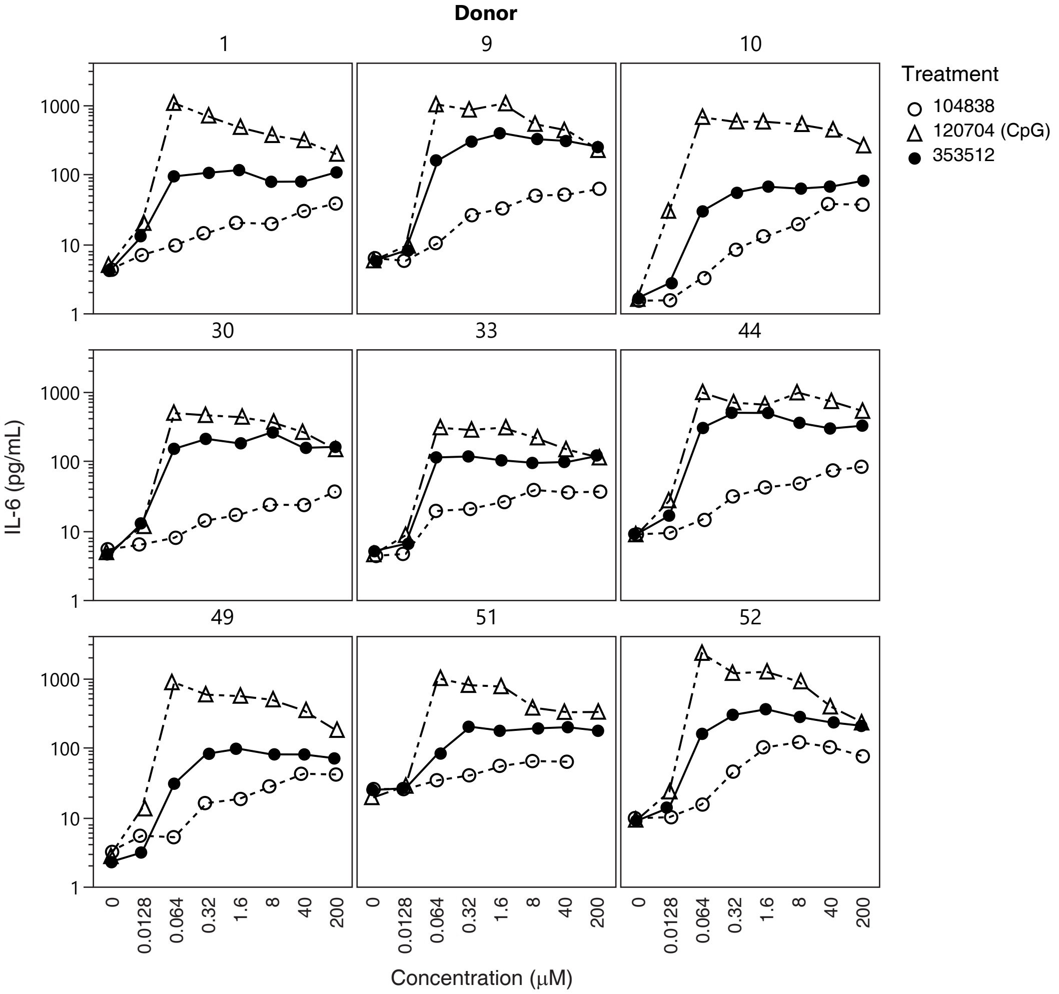


**B**

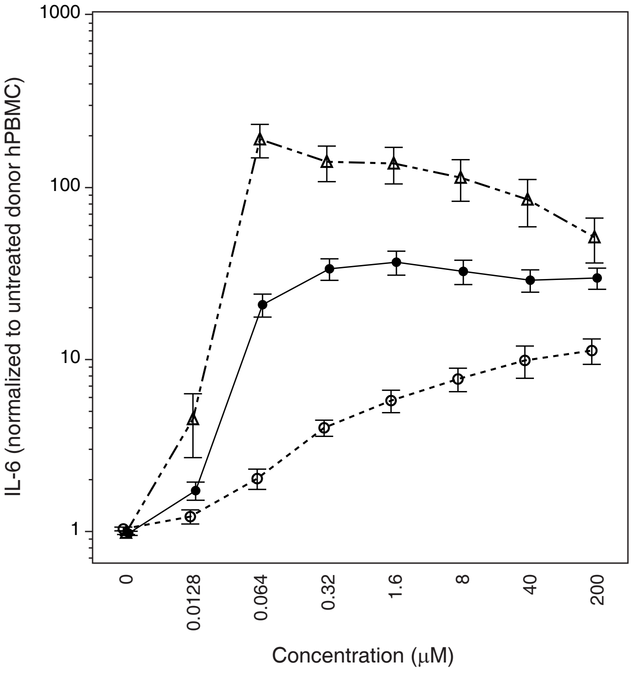
