## Supplemental table 1 for "Early-Stage Identification and Avoidance of Antisense Oligonucleotides Causing Species-Specific Inflammatory Responses in Human Volunteers Peripheral Blood Mononuclear Cells"

| ODN ID | Full Sequence |
|---|---|
| ISIS 104838 | GsCsTsGsAsTsTsAsGsAsGsAsGsAsGsGsTsCsCsC |
| ISIS 113715 | GsCsTsCsCsTsTsCsCsAsCsTsGsAsTsCsCsTsGsC |
| ISIS 120704 (ODN2006) | TsĈsGsTsĈsGsTsTsTsTsGsTsĈsGsTsTsTsTsGsTsĈsGsTsT |
| ISIS 304801 | AsGsCsTsTsCsTsTsGsTsCsCsAsGsCsTsTsTsAsT |
| ISIS 325568 | GsCsAsCsTsTsTsGsTsGsGsTsGsCsCsAsAsGsGsC |
| ISIS 329993 | AsGsCsAsTsAsGsTsTsAsAsCsGsAsGsCsTsCsCsC |
| ISIS 330012 | TsCsCsCsAsTsTsTsCsAsGsGsAsGsAsCsCsTsGsG |
| ISIS 353512 | TsCsCsCsAsTsTsTsCsAsGsGsAsGsAsCsCsTsGsG |
| ISIS 518477 | CsTsCsAsCsAsTsTsGsAsCsAsCsTsGsAsGsG |
| ISIS 818290 (ODN2395) | TsĈsGsTsĈsGsTsTsTsTsĈsGsGsĈsGsĈsGsĈsGsĈsĈsG |
| ISIS 818291 (ODN control) | TsGsĈsTsGsĈsTsTsTsTsGsGsGsGsGsĈsĈsĈsĈsĈsĈ |
